# Decoupling alignment strategy from feature quantification using a standard alignment incidence data structure

**DOI:** 10.1101/2021.02.16.431379

**Authors:** Kwangbom Choi, Matthew J. Vincent, Gary A. Churchill

**Affiliations:** The Jackson Laboratory, 600 Main Street, Bar Harbor, Maine, 04609

**Keywords:** high-throughput sequencing, read alignment, feature quantification, data structure

## Abstract

**Summary:** The abundance of genomic feature such as gene expression is often estimated from observed total number of alignment incidences in the targeted genome regions. We introduce a generic data structure and associated file format for alignment incidence data so that method developers can create novel pipelines comprising models, each optimal for read alignment, post-alignment QC, and quantification across multiple sequencing modalities.

**Availability and Implementation:** `alntools` software is freely available at https://github.com/churchill-lab/alntools under MIT license.

## Introduction

Short read sequencing technologies and their supporting analysis methods represent a core area of bioinformatics research. Due to the variety of experimental strategies used and the scale of the data that can be generated, there is a need for efficient, re-usable data structures that can serve as a link between software tools that generate sequence alignments and downstream analysis tools. An ideal data structure should capture the complete and sufficient information in the alignment output while at the same time eliminating much of the ancillary information. The data structure should be intuitive, easy to incorporate into new software development, and flexible enough to support the most widely used sequencing methods as well as anticipating yet to be developed technologies.

Alignment of short sequence reads to a genome or a set of genomic features is a standard first step in the quantification of sequence data. A variety of alignment strategies are used, from dynamic programming to k-mer based methods, but in most applications the objective is determine how many reads were derived from each of a set of genomic features. Quantification of features would be trivial if each sequenced read aligned uniquely and unambiguously to a single feature. However, features can share regions of high similarity. For example, allelic copies of a gene may differ by only a few single nucleotide polymorphisms (SNPs) or may even be identical. Strategies for dealing with alignment ambiguity range from discarding multiple aligned reads (multi-reads) to sophisticated iterative strategies for weighted allocation of read counts^1;2;3^.

Examples of alignment targets that are composed of discrete genome features include transcripts for aligning RNA-Seq reads, capture regions for whole exome sequencing (WES), digitally digested restriction fragments for reduced representation bisulfite sequencing (RRBS)^4^, and peak regions for chromatin immune precipitation sequencing (Chip-Seq)^5^ or transposase-accessible chromatin sequencing (ATAC-Seq)^6^. The exact location of reads within the feature is often not important as only the number of reads derived from within the feature is needed for quantification. In this case, alignment data can be reduced to an incidence profile that indicates which feature(s) a read aligns to. An incidence profile can be thought of as a binary vector indexed by features with “one” indicating that a read aligns to the feature. Multi-reads are represented by incidence profiles with multiple “ones”. Many reads will have matching incidence profiles and it is sufficient to store only a single profile and the count of the number of reads that share the same profile. The collection of incidence profiles for a sequenced sample or cell is sufficient for feature quantification and provides a substantial data reduction.

Humans and most model organisms have diploid genomes. Allele-specific analysis of sequence data is becoming more common due to longer read platforms and the ability to infer phased gene sequences^7^. An allele-specific alignment target will include two (or more) copies of each feature incorporating their distinct polymorphisms. In applications using multiparent population^8^, features may be represented by multiple allelic types. Allele-specific alignment targets can be generated using `g2gtools` (http://churchill-lab.github.io/g2gtools/) from a reference genome sequence and a variant call format (VCF) file. Interesting applications of allele-specific sequence analysis include demultiplexing pooled libraries of genetically distinct samples^9^ and genotyping by RNA-Seq^10^. In any application that uses an allele-specific alignment target, there will be a high proportion of multi-reads with complex incidence profiles.

There are only few data formats in current use for capturing feature-level read counts. One basic but flexible data format is Loom (http://loompy.org), which can store feature abundance data together with sample and feature meta-data in a single file. Loom can support allele-specific abundance data in multiple layered matrices. However, Loom is not designed to encode alignment incidence profiles. To the best of our knowledge, the only data format that supports alignment incidence profiles is the Barcode-UMI-Set (BUS) file^11^. The BUS file format supports fast quantification and differential analysis in pipelines that utilize `kallisto` and `sleuth` software. Moreover, BUS is specialized for single-cell RNA-Seq data with unique molecular identifier (UMI) barcodes. A BUS file can be generated using the `kallisto` pseudo-aligner and has not been adapted for use with other alignment software. Thus, there is a need for a generic data structure that can support a wide variety of sequencing technologies and is compatible with standard aligners that output binary alignment map (BAM) format files.

We have developed a new file format, ALN, for storing alignment incidence data and have implemented python methods, `alntools`, that perform useful operations on ALN format files. The `bam2ec` method processes BAM files into ALN format. We assume that any post-alignment quality control (QC) operations such as UMI correction or duplicate removal have been performed before conversion because the ALN format captures only the alignment incidence profile data that is needed for feature quantification. The ALN file can be loaded in python as a sparse data object and is accessible to many generic python tools. Our goal is to decouple alignment strategy and post-alignment QC from quantification and downstream analysis to support modular pipeline development. The `alntools` software is freely available under MIT licensing at https://churchill-lab.github.io/alntools/.

## Materials and Methods

The ALN format is composed of a header, an alignment incidence matrix (A), and a count matrix (N). The header captures metadata for samples or single cells (e.g., cell barcodes), as well as feature sequences and metadata including haplotype labels to support allele-specific applications. The A matrix represents alignment incidence profiles for all reads across all samples. The A matrix collapses individual reads into feature equivalence classes — sets of reads that share the same feature mapping pattern. The patterns are stored as a two-dimensional sparse indicator matrix with dimensions of |feature equivalence classes| × |features|. If allele-specific features are used as targets, entries of the A matrix are binary numbers of length |haplotypes| in which each bit represents an alignment incidence on a haplotype (Figure 1a). These binary numbers are converted to decimals when stored in an ALN file. The N matrix is a two-dimensional (| feature equivalence classes| × |samples|) sparse matrix that represents the number of reads or UMIs associated with each feature equivalence class in each sample or cell. Both A and N are stored in Compressed Sparse Column (CSC) format^12^.

**Figure 1:**
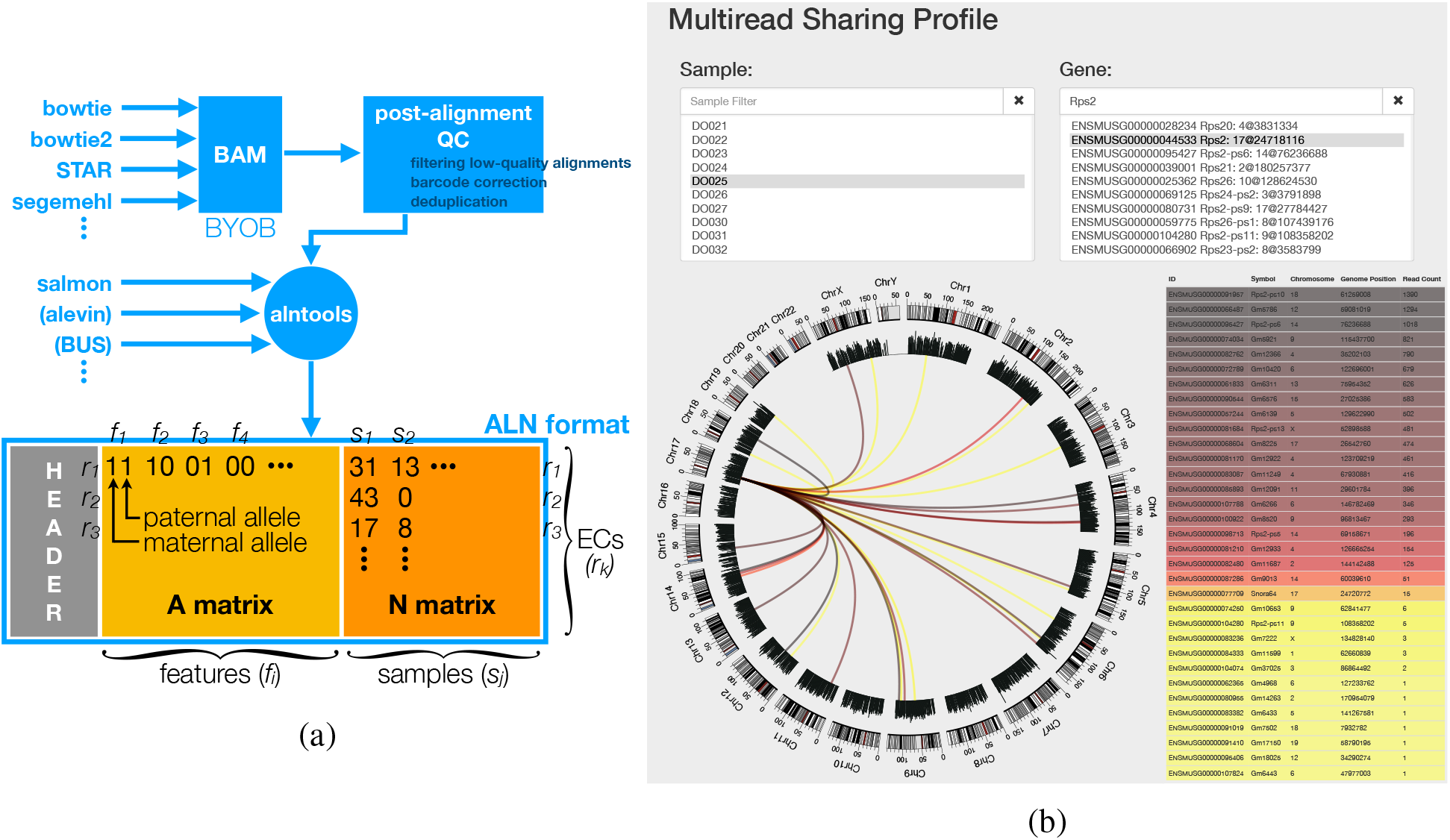
(a) ALN format description. The header contains metadata for feature sequences (*f_i_*) or samples (*s_j_*). The A matrix describes alignment incidence across features for each read or equivalence class (*r_k_*). This matrix can encode allele-specific alignments. For example, 00, 01, 10, and 11 can specify whether r_k_ aligns to the maternal and/or paternal allele of *f_i_*. These binary numbers are converted to decimal — 0, 1, 2, or 3 — when we save it in an ALN format file. The N matrix describes read or UMI counts for each *r_k_* across samples. BUS and `alevin` will be supported in the next version of `alntools`. (b) Alignment incidence data is useful in quality checking for the downstream results besides estimating feature abundance. For example, we can examine whether co-localizing genes for an eQTL is due to sharing multi-mapping reads. We provide an interactive tool that visualizes how a query gene shares multi-mapping reads with other genes.

We have implemented python methods that perform useful operations on ALN files. For example, `alntools` subcommand ‘`bundle`’ collapses transcript alignment incidence of RNA-Seq reads to gene-level incidence; ‘`get_unique_reads`’ identifies uniquely mapping reads or equivalence classes; ‘`view`’ presents how multi-mapping reads are shared among features; or ‘`apply_genotypes`’ filters out alignments inconsistent to genotype calls^10^.

## Results

We created a BAM file with single-cell ATAC-Seq data from a sample of three mouse strains — C57BL/6J, NZO/HlLtJ, and CAST/EiJ — multiplexed for sequencing. We aligned the reads with segemehl^13^ to the common candidate peak regions (or *peakome)* of the three strains by lifting over the reference peakome previously identified from genome alignment. This allele-specific alignment strategy is suitable for simultaneous analysis of demultiplexing and quantification. Processing the BAM file of 65 GB takes 13 minutes to convert into an ALN format of 452 MB on a linux system with 2.2 GHz Intel(R) Xeon(R) E5-2698 v4 (40 threads) and 256 GB RAM. We find that an ALN format file is generally 100-500 times smaller in size than the input BAM file. ALN format does not encode the alignment position or quality sequences.

The utility of ALN format and `alntools`, is best demonstrated by their applications. We implemented an interactive viewer of multi-read sharing among genes (Figure 1b) using `alntools`. The application can be used to visualize sequence features that share multi-mapping reads. We have found this tool to be useful for identifying false positive signals in eQTL analysis that can occur when multi-mapping read counts are not correctly allocated. We also implemented a new version of the `EMASE` algorithm^3^ in C++ that uses ALN format files (http://churchill-lab.github.io/emase-zero/). The new `EMASE` has random access to any chunk of samples in the ALN file which supports execution across multiple cores in a parallel computing environment.

## References

[1] Nicolas L Bray, Harold Pimentel, Páll Melsted, and Lior Pachter. Near-optimal probabilistic rna-seq quantification. Nature Biotechnology, 34(5):525–527, 2016.

[2] Rob Patro, Geet Duggal, Michael I Love, Rafael A Irizarry, and Carl Kingsford. Salmon provides fast and bias-aware quantification of transcript expression. Nature methods, 14(4):417–419, 2017.

[3] Narayanan Raghupathy, Kwangbom Choi, Matthew J Vincent, Glen L Beane, Keith S Sheppard, Steven C Munger, Ron Korstanje, Fernando Pardo-Manual de Villena, and Gary A Churchill. Hierarchical analysis of RNA-seq reads improves the accuracy of allele-specific expression. Bioinformatics, 34(13):2177–2184, 02 2018.

[4] Saurabh Baheti, Rahul Kanwar, Meike Goelzenleuchter, Jean-Pierre A. Kocher, Andreas S. Beutler, and Zhifu Sun. Targeted alignment and end repair elimination increase alignment and methylation measure accuracy for reduced representation bisulfite sequencing data. BMC Genomics, 17(1):149, 2016.

[5] Christopher L. Baker, Shimpei Kajita, Michael Walker, Ruth L. Saxl, Narayanan Raghupathy, Kwangbom Choi, Petko M. Petkov, and Kenneth Paigen. Prdm9 drives evolutionary erosion of hotspots in musmusculus through haplotype-specific initiation of meiotic recombination. PLOS Genetics, 11(1):1–16, 01 2015.

[6] Tang M Giansanti V and Cittaro D. Fast analysis of scatac-seq data using a predefined set of genomic regions. F1000Research, 9:199, 2020.

[7] Olivier Delaneau, Jean-François Zagury, Matthew R. Robinson, Jonathan L. Marchini, and Emmanouil T. Dermitzakis. Accurate, scalable and integrative haplotype estimation. Nature Communications, 10(1):5436, 2019.

[8] Dirk-Jan de Koning and Lauren M. McIntyre. Back to the future: Multiparent populations provide the key to unlocking the genetic basis of complex traits. G3: Genes\Genomes\Genetics, 7(6):1617, 06 2017.

[9] Hyun Min Kang, Meena Subramaniam, Sasha Targ, Michelle Nguyen, Lenka Maliskova, Elizabeth McCarthy, Eunice Wan, Simon Wong, Lauren Byrnes, Cristina M Lanata, Rachel E Gate, Sara Mostafavi, Alexander Marson, Noah Zaitlen, Lindsey A Criswell, and Chun Jimmie Ye. Multiplexed droplet single-cell rna-sequencing using natural genetic variation. Nature Biotechnology, 36(1):89–94, 2018.

[10] Kwangbom Choi, Hao He, Daniel M. Gatti, Vivek M. Philip, Narayanan Raghupathy, Isabela Gerdes Gyuricza, Steven C. Munger, Elissa J. Chesler, and Gary A. Churchill. Genotype-free individual genome reconstruction of multiparental population models by rna sequencing data. bioRxiv, 2020.

[11] Páll Melsted, Vasilis Ntranos, and Lior Pachter. The barcode, UMI, set format and BUStools. Bioinformatics, 35(21):4472–4473, 05 2019.

[12] Iain S Duff, Albert M Erisman, and John K Reid. Direct Methods for Sparse Matrices. Oxford University Press, Inc., USA, 1986.

[13] Christian Otto, Peter F. Stadler, and Steve Hoffmann. Lacking alignments? The nextgeneration sequencing mapper segemehl revisited. Bioinformatics, 30(13):1837–1843, 03 2014.

